# TNF-α induces VE-cadherin-dependent gap/JAIL cycling through an intermediate state essential for neutrophil transmigration

**DOI:** 10.1101/2025.07.20.665739

**Authors:** Jan Philip Kipcke, Maria Odenthal-Schnittler, Mohammed Aldirawi, Jonas Franz, Vesna Bojovic, Jochen Seebach, Hans Schnittler

**Author notes:** Correspondence: Hans Schnittler, Max Planck Institute for Molecular Biomedicine, Röntgenstrasse 20, 48149 Münster, Germany.

## Abstract

Inflammatory endothelial phenotypes describe distinct cellular patterns essential for controlling transendothelial migration of leukocytes (TEM). While TNF-α-induced CAM expression mediates leukocyte interaction, the role of a potential inflammatory morphological phenotype (IMP) – characterised by barrier-function decrease and shape-change in TEM – remains unclear. This study identifies the TNF-α-induced IMP as indispensable for neutrophil TEM, while regulating barrier-function. The TNF-α-induced IMP progresses through two states: an intermediate state that transiently enhances barrier function via MLC-dephosphorylation, junctional actin recruitment and VE-cadherin linearisation, protecting the monolayer from collapse; while the subsequent development of the IMP requires MLC rephosphorylation, junctional actin disassembly, stress fibre formation and Arp2/3-mediated membrane protrusions causing shape-change. This in turn dilutes junctional VE-cadherin, forming intercellular gaps for neutrophil TEM, while inducing junction-associated intermittent lamellipodia (JAIL) to locally restore VE-cadherin adhesion, appearing as gap/JAIL cycles driving junctional dynamics. VE-cadherin overexpression blocks TNF-α-induced IMP and gap/JAIL cycling, reducing TEM by ∼80% without altering CAM expression. These findings highlight gap/JAIL cycling and MLC phosphorylation as key IMP regulators and potential therapeutic targets for inflammatory diseases.

## Introduction

Transendothelial migration (TEM) of neutrophils into inflamed tissue plays a critical role in repair, pathogen defense and wound healing. TEM of leukocytes follows a stepwise cascade of events, characterised by margination/capture, rolling, firm adhesion, spreading and crawling to endothelial exit sites, and ultimately transmigration, also termed diapedesis (1–7). The underlying mechanisms of this process require endothelial activation, mediated by inflammatory mediators (8) or pathogen-associated virulence factors (9) leading to an inflammatory endothelial phenotype that includes the formation of distinct cellular patterns. For example, after TNF-α stimulation, the inflammatory phenotype is characterised by the expression of cell adhesion molecules (CAMs) for interaction with leukocytes, but also shows a change in endothelial morphology and dynamics, accompanied by a downregulation of barrier function here referred to as the inflammatory morphological phenotype (IMP). While CAM expression is essential for the interaction of neutrophils with the endothelium, the role of TNF-α-induced morphological remodelling leading to the IMP remains unclear for TEM.

TEM of neutrophils occurs either through the cell body (transcellular), as in the brain (10–12), or between endothelial cell junctions (paracellular TEM), occurring in most organs (13, 14). TEM is initiated by leukocyte-endothelial interactions via mediator-induced CAMs expressed on the endothelial surface, which serve as receptors for leukocyte adhesion. For example, lymphocyte function-associated antigen 1 (LFA-1) is a heterodimeric integrin, expressed in neutrophils, and interacts with ICAM-1, causing ICAM-1 clustering (2, 5, 15, 16). As a consequence, endothelial actin is recruited to this site, forming a ring-like structure around the transmigrating neutrophil. In addition, membrane protrusions known as ‘docking structures’ or ‘transmigratory cups’ may appear, which are thought to contribute to TEM as well (17–19). However, the molecular mechanisms by which the endothelium contributes to or controls neutrophil transmigration remain debated, although cytoskeletal and junctional proteins are widely recognised as key players in paracellular TEM (6, 20–24) (25). This includes, in addition to the actin cytoskeleton, vascular endothelial (VE) cadherin (26–30), JAMs (31), CD99, as well as PECAM-1 present in the lateral border recycling compartment (LBRC), which may act as a gateway for leukocytes (32, 33).

Endothelial adherens junctions, composed of the VE-cadherin/catenin complex functionally and dynamically linked to actin filaments, are key structures for organising and regulating cell contacts. This includes a specific role for actin filament stress fibres and branched actin filament dynamics, which are differentially regulated by EPLIN isoforms during neutrophil TEM (25). Importantly, antibody-mediated VE-cadherin inactivation (34, 35), and Y-phosphorylation or dissociation of VE-PTP from VE-cadherin increase transendothelial migration (TEM) (36–38), whereas stabilisation of the VE-cadherin-actin interaction via a VE-cadherin-α-catenin chimera reduces leukocyte extravasation (39). Other work has proposed a dissociation of the VE-cadherin/catenin complex from actin filaments. (40–43). These findings are consistent with the widely accepted role of micrometre-sized pores at endothelial junctions lacking VE-cadherin as exit sites during TEM. After TEM, there is also agreement that VE-cadherin adhesion is reconstituted, involving actin-driven membrane protrusions as shown previously (24, 44–46). Although these data clearly highlight the critical role of junctional molecules in TEM, the TNF-α-induced IMP, with upregulation of cell and junctional dynamics, morphological changes and barrier function breakdown, requires further mechanistic understanding for the neutrophil TEM (47–51).

Here, we investigated the functional role of the TNF-α-induced inflammatory morphological phenotype (IMP) in the control of barrier function and neutrophil TEM. This study builds on previous research in angiogenesis and wound healing by focusing on actin– and VE-cadherin-mediated endothelial shape changes and dynamics. Specifically, VE-cadherin remodelling and shape change occur through actin-driven junction-associated intermittent lamellipodia (JAIL), controlled by Rac-1-activated Arp2/3-complex, creating branched actin filaments that form at sites of reduced VE-cadherin to restore adhesion (52–56). Indeed, we establish that the fully developed TNF-α-induced inflammatory endothelial phenotype, in addition to the well-characterised upregulation of cell adhesion molecules (CAMs), is further defined by an inflammatory morphological phenotype (IMP) involving intercellular gap formation and actin-driven junction-associated intermittent lamellipodia (JAIL). The alternating formation of intercellular gaps and JAIL generates gap/JAIL cycles, which we demonstrate regulate VE-cadherin-mediated endothelial barrier function during inflammation and are indispensable for neutrophil TEM.

## Materials and Methods

Reagent and Resource Table.

Reagent and Resource Table
| Reagent or Resource | Source | Identifier number |
| --- | --- | --- |
| **Antibodies and Chemicals Immunofluorescence** |  |  |
| Goat anti-VE-Cadherin | R&D (Minneapolis, USA) | Cat.#AF938 RRID:AB_355726 |
| Goat anti-ICAM-1 | Santa Cruz (Dallas, USA) | Cat#sc-8439<br>RRID:AB_2905502 |
| Mouse anti-Beta-Catenin | (BD) Transduction<br>(Hamburg, Germany) | Cat#610154 RRID:AB_397555 |
| Phalloidin-TRITC | Sigma-Aldrich (St. Louis, USA) | Cat#P1951; RRID:AB_2315148 |
| Phalloidin-TRITC 647 | Abcam (Cambridge, UK) | Cat#ab176759 |
| DAPI | Roche (Penzberg, Germany) | Cat#10236276001 |
| Alexa 488 donkey anti-goat | Invitrogen (Carlsbad, USA) | Cat#A11055;<br>RRID:AB_2534102 |
| Alexa 568 donkey anti-mouse | Invitrogen (Carlsbad, USA) | Cat#A10037<br>RRID:AB_2534013 |
| Alexa 633 donkey anti-goat | Invitrogen (Carlsbad, USA) | Cat#A21082<br>RRID:AB_141493) |
| Alexa 647 donkey anti-mouse | Invitrogen (Carlsbad, USA) | Cat#A31571<br>RRID:AB_162542) |
| **Antibodies Western Blot** |  |  |
| Goat anti-VE-Cadherin | R&D (Minneapolis, USA) | Cat.#AF938 RRID:AB_355726 |
| Mouse anti-GFP/EGFP | Thermo Fisher Scientific (Waltham, USA) | **Cat#MA5-15256**<br>RRID:AB_10979281 |
| Mouse anti-ICAM-1 | R&D (Minneapolis, USA) | Cat#BBA3<br>RRID:AB_356950 |
| Mouse anti-Alpha Tubulin T5168 (Clon B-5-1-2) | Sigma-Aldrich (St. Louis, USA) | Cat#T5168 RRID:AB_477579 |
| Mouse anti-beta-Catenin | Santa Cruz (Dallas, USA) | Cat#sc-7199 RRID:AB_634603 |
| Rabbit anti-EGFP | Abcam (Cambridge, UK) | Cat#ab6556<br>RRID:AB_305564) |
| IRDye® 680RD donkey anti-goat | LICOR Bioscience (Lincoln, USA) | Cat#926-32224;<br>RRID:AB_1575081 |
| IRDye® 680RD donkey anti-mouse | LICOR Bioscience (Lincoln, USA) | Cat#926-32222;<br>RRID:AB_621844 |
| IRDye® 800RD donkey anti-goat | LICOR Bioscience (Lincoln, USA) | Cat#926-32214;<br>RRID:AB_621846 |
| IRDye® 800RD donkey anti-mouse | LICOR Bioscience (Lincoln, USA) | Cat#926-32212;<br>RRID:AB_621847 |
| IRDye® 800 RD donkey anti-rabbit | LICOR Bioscience (Lincoln, USA) | Cat#926-32213;<br>RRID:AB_621848 |
| **Antibodies MLC-Phosphorylation Analysis** |  |  |
| Rabbit anti-human-pmlc2-serin19 | Cell Signaling | Cat#3671 |
| Mouse anti-myosin | Sigma | M4410 |
| IRDye® 680RD Goat anti-Mouse IgG | LICOR Bioscience (Lincoln, USA) | Cat#926-68070 |
| IRDye® 800CW Goat anti-Rabbit IgG | LICOR Bioscience (Lincoln, USA) | Cat#926-32211 |
| **Reagents and Materials Flow Assay** |  |  |
| HBSS 10x with Magnesium and Calcium | Thermo Fisher Scientific (Waltham, USA) | Cat#14065056 |
| HEPES solution | Sigma-Aldrich (St. Louis, USA) | Cat#H0887 |
| Sodium Bicarbonat 7,5% | Thermo Fisher Scientific (Waltham, USA) | Cat#25080094 |
| ACK Lysis Buffer | Thermo Fisher Scientific (Waltham, USA) | Cat#A1049201 |
| Histopaque 1077 | Sigma-Aldrich (St. Louis, USA) | Cat#10771 |
| Histopaque 1119 | Sigma-Aldrich (St. Louis, USA) | Cat#11191 |
| **Bacterial and Virus Stains** |  |  |
| EGFP, Lentivirus | EGFP, Lentivirus<br>Hofer et al., 2018 | N/A |
| EGFP-p20, Lentivirus | Abu Taha et al., 2014 | N/A |
| VE-cadherin-mCherry, Lentivirus | Abu Taha et al., 2014 | N/A |
| Life-Act-EGFP, Lentivirus | Abu Taha et al., 2014 |  |
| VE-cadherin-EGFP, Adenovirus | Kametami & Takeichi, 2007 | Kind gift |
| **Experimental Models: Cell Lines** |  |  |
| Human: umbilical cord vein endothelial cells (HUVEC) | This paper | N/A |
| Polymorphonuclear leukocytes | This paper | N/A |
| **Software and Algorithms** |  |  |
| Cell Border Tracker | Seebach et al., 2015 | N/A |
| Fiji | N/A | https://imagej.net/software/fiji/ |
| MATLAB | Mathworks | N/A |
| Zen | Carl Zeiss Jena | N/A |
| Odyssey | LICOR | N/A |
| Chemotaxis and Migration Tool | IBIDI | N/A |
| Good Notes Application | Apple Store | N/A |

## Lead Contact and Materials Availability

Further information and requests for resources and reagents should be directed to and will be fulfilled by the Lead Contact, Hans Schnittler. All unique/stable reagents generated in this study are available from the Lead Contact with a completed Materials Transfer Agreement.

## Cell culture and coatings

Human umbilical vein endothelial cells (HUVEC) were isolated and cultured in endothelial cell growth medium (Promocell, Heidelberg, Germany), following procedures outlined previously (57). The utilization of HUVEC adhered to ethical principles outlined in the Declaration of Helsinki and received approval from the ethics committees at University of Muenster (2009-537-f-S). For experiments, HUVEC from passage 0 were seeded on crosslinked gelatine coated culture supports (57). All used culture supports were coated prior to cell seeding with 0.5% gelatine/PBS for 30 min at 37 °C, crosslinked with 2% glutaraldehyde for 10-15 min, and incubated in 70% ethanol/water for 30-60 min at room temperature (RT). After five PBS washes, dishes underwent overnight exposure to 2 mM glycine/PBS at RT. This coating allows the deposition of extracellular matrix proteins by the cells, contributing to a more natural cellular behavior (58). Before cell seeding, culture dishes underwent five consecutive PBS washes.

Polymorphonuclear neutrophils from human blood were isolated through density gradient centrifugation (Histopaque®-1077, Histopaque®-1119) after obtaining formal consent from healthy donors.

## DNA constructs and viral transductions

Recombinant lentiviral vectors carrying EGFP-p20, LifeAct-EGFP and VE-cadherin-mCherry have previously been employed (52, 53). Adenovirus carrying VE-cadherin-EGFP and EGFP were kindly provided by Masatoshi Takeichi (59).

## Lentivirus production

Lentivirus particles were generated in HEK 293T cells cultured in DMEM (high glucose) medium in 15-cm cell culture dishes. Transfection involved the pFUGW vector carrying the gene of interest, along with the packaging vectors (pCMV-ΔR8.74) and the VSV glycoprotein-carrying vector (pMD2G). Specifically, plasmids pFUGW-gene of interest (23 µg), pCMV-ΔR8.74 (23 µg), and pMD2G (11.5 µg) were dissolved in 1725 µl DMEM medium w/o FCS and antibiotics (DMEM−/−) (solution A). Simultaneously, the transfection solution, consisting of 172 µl of the transfection reagent PEI (1 mg ml−1), was dissolved in 1600.8 µl DMEM (−/−) (solution B). Both solutions were incubated at RT for 30 min and an additional 30 min once mixed. The mixture was added dropwise to the 293T cells. After 14–16 h of incubation, a medium exchange was performed using 20 ml fresh, pre-warmed DMEM supplemented with 10% FCS, 1% Pen/Strep, and 1% Glutamine. After 24 h of culturing, the medium was collected and cleared at 3000 rpm for 10 min. The supernatant was filtered through 0.45-µm filters, and the virus was concentrated by ultracentrifugation (1.5 h at 25,000 rpm; 4 °C). The pellet was resuspended in 150 µl PBS containing 1% BSA, and aliquots were stored at −80 °C until use.

## Adenovirus amplification

Adenoviruses carrying VE-cadherin-EGFP or EGFP were amplified by the following steps: HEK 293 cells were seeded in T75 cm^2^ tissue culture flasks and cultured until reaching 80– 90% confluency after 24 h. The medium was changed to DMEM without FCS and P/S supplemented with 1% Glutamine and 10 µl (15 MOL) of stock adenovirus. After 1 hour of incubation, equivalent volumes of DMEM supplemented with 4% FCS and 1% Glutamine was added to reach a final concentration of 2% FCS in culture medium. After an additional 2 days of culturing, the virus was released by four cycles of freeze-thaw-vortex. To increase the virus yield, additional culture cycles can be performed by distributing the respective suspension to 5 or 10 times greater HEK293 cell culture area (80–90% confluency), culturing for two days, and processing with another four cycles of freeze-thaw-vortex. For virus purification, the cells were scraped from the substrate and centrifuged at 500 × g for 10 min.

The cell pellet was resuspended in 6 ml sterile PBS followed by another four cycles of freeze-thaw-vortex. The lysate was cleared by 4000 x g at 4 °C for 20 min.

For further purification, 6 ml of the supernatant was mixed with 3,3 gr ultrapure CsCl, vortexed, and separated at 1,76 × 105 × g at 10 °C for 22 h. The virus fraction was collected with glycerol (10% final concentration), dialyzed against BSS, aliquoted and stored at −80 °C until use.

## Lentivirus– and adenovirus-mediated gene transductions

For lentivirus transductions, HUVEC were incubated with viral particles suspended in endothelial growth medium (-/-) supplemented with 3% poly-vinyl-pyrrolidone (PVP) for 1 hour at 37°C to enhance transduction efficiency. Endothelial growth medium (+/+) was then added, and cells were cultured further before being subjected to experiments after 24–72 h. Protein expression of interest was assessed using immunofluorescence and Western blotting. For adenovirus transductions, HUVEC were incubated with viral particles in endothelial growth medium (-/-) containing 3% PVP for 4 h at 37°C. The medium was subsequently changed to endothelial growth medium (+/+) and cells were further incubated for 16 h before being used in experiments.

## Western blot analysis

For western blot analyses, HUVEC cultures were lysed in 6% sample buffer, as described previously (53). The protein concentration of the cell lysates was determined using the Amido-black protein assay (60). Western blotting was carried out following standard protocols, with the same amounts of protein loaded onto each lane (60). After protein transfer via wet blotting, membranes were incubated with appropriate antibodies and subsequently detected using secondary near-infrared-labelled antibodies with the Li-Cor system (Li-Cor Odyssey Infrared Reading System, Homburg, Germany). Quantitative analyses were performed using the Odyssey software package (Li-Cor).

## Myosin light chain phosphorylation analysis

Phosphorylation of myosin light chains at serin 19 was determined using the In-Cell Western Assay (LICOR). HUVEC P1 were seeded at 12 000 cells/well in a 96-well plate (Greiner, REF: 655090) in endothelial growth medium as described above. Prior to seeding, cultures dishes were coated with crosslinked gelatine (see cell culture and coatings). After 4 days in culture, the cells were treated with 50 ng/ml of TNF-α for time periods as indicated in the experiments. Subsequently, cells were fixed with 4% paraformaldehyde for 10 min, permeabilized with 0.1% Triton X-100, and blocked with 5% donkey serum. Wells for background subtraction were treated following the In-Cell Western protocol. For In-Cell Western analysis of myosin light chain phosphorylation, rabbit anti-human-pmlc2-serin19 and mouse anti-myosin were used as primary antibodies. IRDye® 680RD Goat anti-Mouse IgG and IRDye® 800CW Goat anti-Rabbit IgG from LICOR were used as secondary antibodies. Plates were imaged at 700nm and 800nm using the LICOR Odyssey CLX Imaging System. The fluorescent intensity of phosphorylated MLC was normalized to the fluorescent intensity of total myosin light chain on the same well after background subtraction using Empiria Studio Software (LICOR).

## Immune labelling of HUVEC

For immunolabelling of HUVEC proteins, cells were fixed with freshly prepared 4% paraformaldehyde supplemented with 0.88mM of calcium dissolved in PBS for 10 min at room temperature. After fixation, cells were washed three times for 5 min each using PBS with 1% BSA and 0.88mM of calcium (PBS/BSA). Subsequently, cell membranes were permeabilized with 0.1% Triton X-100 in PBS for 10 min at room temperature, followed by three additional washes with PBS/BSA 1%. Next, cells were incubated with the respective primary antibody overnight at 4°C, washed again, and exposed to appropriate secondary antibodies. Finally, cultures were mounted in Dako fluorescence mounting medium and evaluated using either LSM 780 (Zeiss), or SIM.

## Quantification of morphology

For phase-contrast-based quantification of the aspect ratio, HUVEC were imaged for 6 h after TNF-a stimulation with 10x magnification (1800 +/-200 cells). Morphometric analysis was performed using Matlab and Fiji/ImageJ (Version 1.49p) (61) 2012) using custom-made codes. For quantification of the HUVEC perimeter, LSM images of HUVEC labelled for VE-cadherin were first segmented using our self-developed software Cell Border Tracker (CBT) (62), and then the perimeter was determined using Fiji/Image J.

## Quantification of dynamics, relative protein concentrations and line scans of actin

Quantification of cell migration was performed by manual cell tracking from time lapse recordings using the Fiji ‘Manual Tracking’ plugin and the ‘Chemotaxis and Migration Tool’. Migration was analysed by comparing parameters, including i) accumulated distance (the total distance travelled by a cell) and ii) velocity. Relative β-catenin and actin levels in cell cultures were quantified using CBT software, as described (62). For the measurement, the integrated protein intensity of the region of interest (ROI) within 3-5 pixels adjacent to the cell border was divided by the total cell border length of the ROI. Line scans of the junctional actin intensity were performed orthogonally to the cell border using Fiji. Maxima of the actin intensity were considered for each time point.

## Comparative demonstration of cell displacements in LifeAct expressing HUVEC monolayer and kymographs of cell junction dynamics

Analyses were performed from time-lapse recorded LifeAct-EGFP expressing HUVEC using Fidji. For grey scale difference analysis, images were converted to 8-bit grey scale, followed by Multi Kymographs stack differences. After selecting the region of interest (ROI), multiple measures were applied. Kymographs were generated after converting image stacks to 8-bit greyscale. Selected lines were analysed by kymograph builder plaque in.

## Transendothelial electrical resistance (TER) measurements during TNF-a application

TER measurements of HUVEC under TNF-a treatment were performed at a shear stress of 1 dyn/cm^2^ using the cone-and-plate based BioTech-Flow system (MOS Technologies, Telgte, Germany), as described in detail previously (63).

## Microscopy

For phase-contrast live-cell imaging, cells were seeded on a crosslinked gelatine-coated glass-bottom dishes (Maedler-Chambers, (MOS-Technologies,Germany) and Willco Wells (Amsterdam/Netherlands)) and automatically imaged using an Axio observer Z1 (Carl Zeiss, Oberkochen, Germany) supplied with water-vapour-saturated 5% CO2/air and a temperature of 37°C using an LDA-plan 10× objective lens.

For fluorescence live-cell imaging, cells were seeded on crosslinked gelatine-coated glass-bottomed dishes as described above. Time-lapse imaging was performed using an Axio observer Z1 confocal microscope equipped with Yokogawa Spinning Disc Unit (SDM), (Carl Zeiss, Oberkochen, Germany) and definite focus. Cells were inserted in a 37°C heated stage providing water-vapour-saturated 5% CO2/air. Live cells were imaged using a heated Plan-A Apochromat 40x/1,4 Oil DIC (UV) VIS-IR objective and the appropriate lasers for excitation and emission. Laser scanning microscopy (LSM) and structural illumination microscopy (SIM) was carried out using a Zeiss Elyra/LSM 780 microscope (Carl Zeiss, Oberkochen, Germany). Immunofluorescent images were acquired using a 63× plan-Apochromat oil immersion objective and an electron-multiplying CCD camera using 405, 488, 563, and 647 nm diode lasers. Images were reconstructed and processed for structure illumination with Zen imaging software (Carl Zeiss, Oberkochen, Germany). Alignment parameters for the different channels were obtained from measurements of a multispectral calibration slide (170 nm multi-wavelength fluorescent beads, Carl Zeiss) taken with the same camera setup as the biological samples.

## Phase contrast leukocyte transmigration assay under flow conditions

Experiments were performed using the self-constructed “Insert Flow Chamber” for leukocyte transmigration assays under flow conditions (patent no. 10 2020 131 894). The chamber is designed so that cells can be cultivated under normal standard conditions, completely avoiding any deficiency of nutrients and oxygen. The chamber provides unidirectional laminar flow as demonstrated by both numerical simulation and dye flow experiments (25). HUVEC were cultured on cross-linked gelatine-coated Willco glass bottom wells using insert flow seeding equipment. After reaching a cell density of 1-1.2x 10^5^ cells/cm^2^, TNF-α stimulation (50ng/ml) was performed for 6 h. The flow through the insert flow chamber was adjusted to 1 dyn/cm^2^ at 37°C and 5% CO_2_/air. 1x 10^6^ freshly isolated neutrophils suspended in HBSS buffer were injected via a side port and time-lapse imaging was performed using the Axio Z1 automated observer (Carl Zeiss, Göttingen, Germany). Four different fields were captured at 10x magnification for 30 min and used for quantitative measurements.

## Quantification of neutrophil adhesion and transmigration on HUVEC monolayers based on phase contrast time lapse recordings

To determine the amount of total interacting PMN, cells either adherent to the endothelium or already transmigrated were counted 3 min after neutrophil injection. Transmigrated neutrophils were distinguished from adherent neutrophils by the transition from bright to dark phase at 3, 6, 9 and 12 min. Transmigration of neutrophils was calculated for each time point as follows: transmigrated neutrophils / total interacting PMN at 3 min. Movies were analysed using the Fiji/Image J software and the Good Notes application.

## Fluorescence live cell imaging of leukocyte transmigration under flow conditions

The self-constructed flow chamber was also used for live cell imaging. HUVEC were transduced with lentiviruses (EGFP-p20, VE-cadherin-m-Cherry and Life-Act-EGFP) or adenoviruses (VE-Cadherin-EGFP) carrying the protein of interest. After 6 h of TNF-a stimulation (50ng/ml), 1x 10^6^ freshly isolated PMN suspended in HBSS buffer were drawn onto HUVEC monolayers seeded in flow chambers at 1 dyn/cm^2^ at 37°C. Images were captured using a spinning disc confocal microscope (SDM (Carl Zeiss, Göttingen, Germany)).

## Statistics

Statistical analyses were performed using GraphPad Prism (GraphPad Software, San Diego, CA, USA). If the data followed a Gaussian distribution, comparisons between two groups were conducted using Student’s *t*-test, while comparisons among more than two groups were performed using one-way ANOVA. If the data did not follow a Gaussian distribution, Mann-Whitney *U* test was used for two-group comparisons, and Kruskal-Wallis test was applied for comparisons involving more than two groups. A *p*-value of less than 0.05 (*p< 0.05, **p< 0.01, *** p< 0.001, **** p< 0.0001) was considered statistically significant. Data are presented as mean ± SEM.

## Results

### TNF-α induces an actin-based intermediate state before switching to an inflammatory morphological phenotype

The pro-inflammatory cytokine TNF-α is known to upregulate the expression of adhesion molecules (CAMs), thereby facilitating endothelial-leukocyte interactions. In addition to these effects, TNF-α induces morphological changes in endothelial cells associated with reduced barrier function, here referred to as the inflammatory morphological phenotype (IMP). This phenotype is often considered to represent endothelial dysfunction (8, 49, 51, 64, 65). However, it remains questionable whether TNF-α-induced cell and junctional dynamics are regulated despite inflammation and whether this inflammatory morphological phenotype (IMP) contributes to neutrophil TEM.

To investigate and characterise the time course and mechanisms underlying the development of IMP, we used HUVEC cultures (1-1.2 x 10⁵ cells/cm²) as a model, representing quiescent endothelium with well-established adherens junctions. Treatment of confluent HUVEC with 50 ng/ml TNF-α converted them to an inflammatory phenotype within hours, characterised by ICAM-1 expression (Supplementary Fig. S1a, S1b) and a time-dependent change in cell shape with increased perimeter and aspect ratio, as determined from VE-cadherin immune labelling (Figure 1a; Supplementary Fig. S1c, d; Movie 1). These morphological changes were accompanied by enhanced migration speed and accumulated distance travelled within the cell monolayer, as analysed by time-lapse-imaging (Figure 1b,1c; Supplementary Fig. S1e). In addition, we investigated the change in TNF-α-induced barrier function in the presence of shear stress at 1 dyn/cm^2^ to mimic the conditions used for neutrophil TEM. Barrier function was monitored by determination of transendothelial electrical resistance (TER) from impedance spectroscopy measurements. Intriguingly, shortly after TNF-α application, there was a rapid and transient increase in TER of ∼20% within the first hour (Figure 1d), representing enhanced endothelial barrier integrity. Although similar transient increases in TER have been noted in prior studies, even in the absence of shear stress (57, 66–68), they have not been explicitly analysed or discussed.

**Figure 1.**
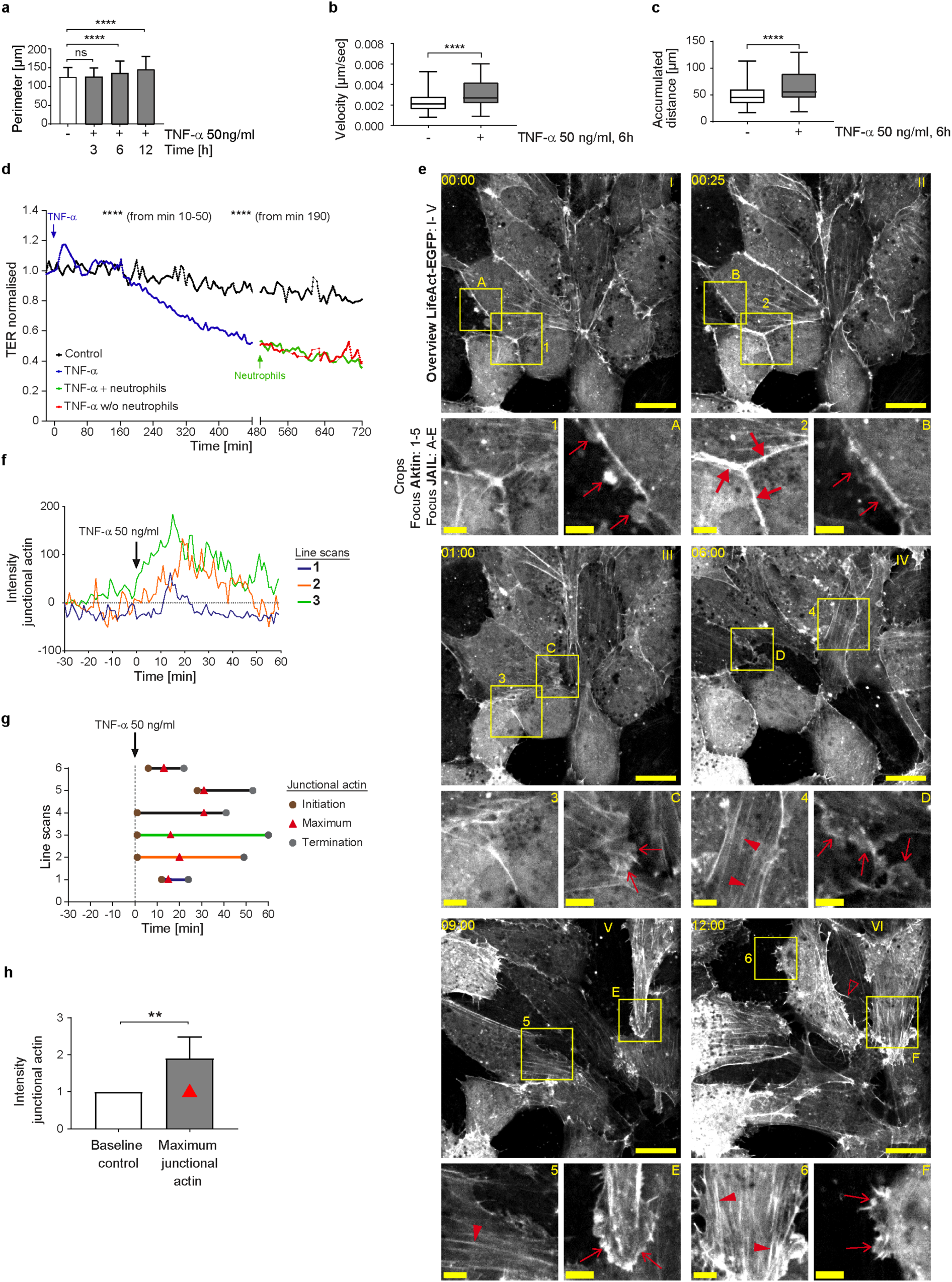
TNF-α-induced endothelial inflammatory pathotype is triggered via a cell-protective, actin-based intermediate state. **a-c)** Time-dependent analyses of morphodynamic parameters in HUVEC cultures (1-1,2 x 10^5^ cells/cm^2^) upon TNF-α treatment. **a)** Cell perimeter; n= 3 independent experiments with No of cells analysed NT = 569; TNF-α 3h = 1017 cells; TNF-α 6h = 935; TNF-α 12h = 810. Kruskal-Wallis test. **b)** Cell migration velocity and **c)** accumulated distance; n= 3, considering 36 single cells per experiment; Mann-Whitney-U-Test. **d)** Mean values of transendothelial electrical resistance (TER) based on impedance spectroscopy measurements (1 minute interval) in independent experiment. Arrows indicate the addition of TNF-α and neutrophils, respectively. Control, n= 2, 9 electrodes in 4 chambers; plus TNF-α, n=3, 22 electrodes in 8 chambers until time point 480 min; +/-neutrophils n= 3, 14 electrodes in 5 different chambers. **e)** Representative time lapse images of HUVEC cultures (1-1.2 x 10^5^ cells/cm^2^) expressing LifeAct-EGFP to monitor actin dynamics in response to TNF-α stimulation at indicated time points (00:00, hh:mm, scale bar: 20 µm). Cropped areas shown below, scale bar: 5 µm; each time point is indicated by letters and numbers. Large red arrows = junctional actin; small red arrows = actin-driven membrane protrusions; red arrowheads = stress fibres; red unfilled arrowheads = filopodia. Note: TNF-α induced actin recruitment is followed by disassembly together with stress fibre and protrusion formation (n=3 independent experiments). **f-h)** Line scans of junctional actin based on LifeAct-EGFP expression in HUVEC; selected areas are indicated in Supplementary Figure S 1. **f**) Time-dependent 3 representative line scans; arrow = TNF-α application. **g)** Maxima of junctional actin recruitment. **h)** Relative actin recruitment determined by intensity ratio of maximal recruitment divided by actin intensity prior to TNF-α stimulation (Mann-Whitney-U-test). ns=not significant.

To elucidate the structural and molecular basis of this phenomenon, fluorescence live-cell imaging of key junctional molecules – fluorescently labelled actin and VE-cadherin – was performed in response to TNF-α. Under control conditions, confluent HUVEC expressing fluorescently labelled LifeAct exhibited low actin dynamics, sparse stress fibre formation, moderately developed junctional actin and limited actin-driven membrane protrusions (Figure 1e; Movie 2, control). These protrusions were previously identified in endothelial cells as junction-associated intermittent lamellipodia (JAIL), which dynamically maintain endothelial integrity (for review see (56)). Intriguingly, coinciding with the transient TER increase, fluorescent time-lapse analysis revealed a two-state process in actin filament organisation and dynamics after TNF-α administration. Within the first 28 minutes after TNF-α application, prominent, transiently appearing junctional actin was formed (Figure 1e – g; Movie 2), characterised by line scanning at different positions (Supplementary Fig. S1f). The intensity peak of actin recruitment reached approximately two times the baseline level and occurred between 13 – 31 minutes post-stimulation (Figure 1g-h; Movie 2), a timeframe consistent with TER increase (Figure 1d). Based on these early transient functional and morphological changes induced by TNF-α, which contrast with the later inflammatory phenotype, we propose this phenomenon as an intermediate state. Similar observations have been reported in developmental biology, where such intermediate states are thought to coordinate cellular behaviour (69, 70). After this intermediate state, the subsequent second phase, towards a fully IMP, was characterised by a general increase in actin dynamics. Specifically, there was a progressive disassembly of junctional actin and an increase in actin-driven membrane protrusions with few filopodia, together with the sequential development of stress fibres (Figure 1e), as demonstrated by time-lapse imaging. These changes in actin dynamics align with barrier function decrease and upregulation of morphodynamics (Figures 1a-f; Supplementary Fig. S1a-S1e, Movie 2). Ultimately, this process leads to the fully developed inflammatory morphological phenotype of the endothelium.

### Characterisation and mechanistical details of the TNF-α induced intermediate state

Obviously, the intermediate state induced by TNF-α strengthens endothelial integrity and prevents immediate barrier breakdown by promoting actin recruitment to the cell junctions. This intermediate state resembles the mature, quiescent endothelial actin phenotype typically observed in large venous endothelial cells in vivo (71–74). We therefore propose this intermediate state serves to resets the endothelium to a baseline organisational level, potentially initiating a hierarchical cascade that facilitate the controlled remodelling towards an IMP. Consistent with the established role of MLC-phosphorylation as a regulator of barrier function, junctional actin formation, stress fibre assembly and contractility, (75–77), we further found that transient actin recruitment to cell junctions after TNF-α application is accompanied by transient MLC-dephosphorylation within 10-30 minutes (Figure 2a). MLC dephosphorylation-induced tension loss caused moderate cell displacements within 0-60 minutes of TNF-α treatment, as quantified by grey level changes in phase-contrast time-lapse series (Figure 2b). During this time period, actin-driven membrane protrusions were largely arrested (Figure 2c, t= 0-50 min), whereas the control-period kymographs of the cells prior to TNF-α application showed moderate protrusion dynamics at cell junctions (Figure 2c, t= –30-0 min). Correlation analyses revealed two key relationships during the transient intermediate state. A robust negative correlation between myosin light chain (MLC) phosphorylation and junctional actin (Figures 2d, 2e) and a strong positive correlation between junctional actin and TER (Figures 2d, 2f). The correlation between MLC-phosphorylation and TER also tended to be negative (r = 0.7033), but did not reach statistical significance (Figures 2d, 2g). This lack of significance may reflect the temporal precedence of MLC-dephosphorylation relative to TER elevation. These parameters were measured in completely independent experiments, strongly indicating a coordinated cellular response and highlights a potential mechanistic link in the conversion of endothelial cells via an intermediate state towards an inflammatory morphological phenotype.

**Figure 2.**
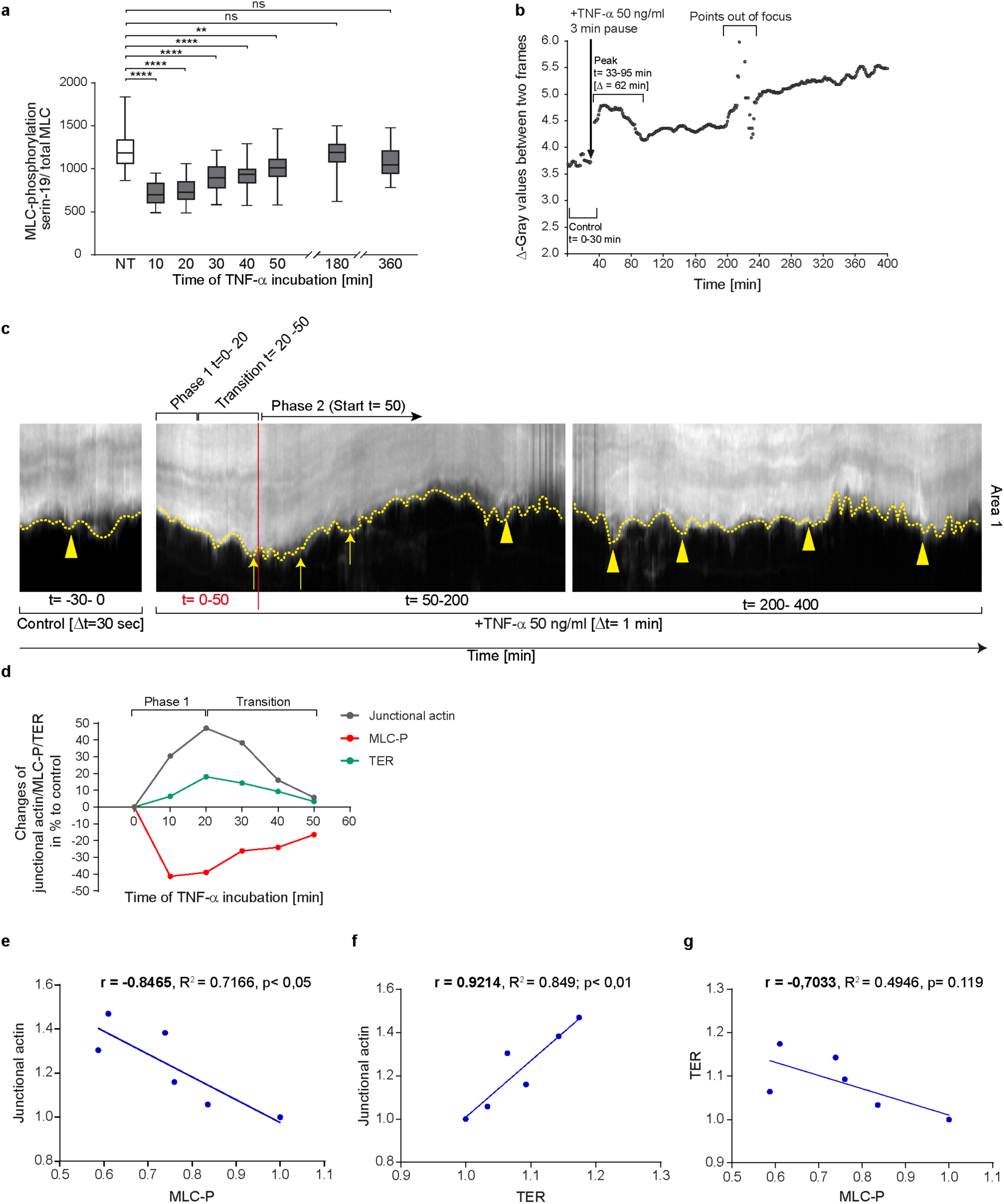
Correlation of MLC-dephosphorylation, actin recruitment and increase in TER, as well as changes in membrane protrusions, characterises the TNF-α-induced intermediate state. **a)** Ratio of serin-19 MLC phosphorylation and total MLC after TNF-α stimulation in confluent HUVEC cultures (1-1.2 x 10^5^ cells/cm^2^) by antibody labelling using *In Cell Western Assay*, (Licor); n= 3 independent experiments with NT, n=20; 10min, n=30; 20min, n=20; 30min, n=30; 40min, n=20; 50min, n=30; 3h, n=30; 6h, n=30. Each n represents 0.34cm^2^ of confluent HUVEC (10^5^ cells/cm^2^). One-way ANOVA. **b)** Change in total monolayer dynamics after TNF-α stimulation analysed by grey level difference analysis from phase-contrast-based time-lapse imaging (image frequency: control, 2 images/min; after TNF-α stimulation 1 image/min). **c)** Representative kymographs of cell junction dynamics in LifeAct-EGFP expressing HUVEC, with treatments and time points as indicated. **d)** Overlay of MLC phosphorylation levels (from data shown in Figure 2a); maximum junctional actin recruitment (from data shown in Figure 1g); TER (from data shown in Figure 1d). **e-g)** Pearson correlation analyses between actin, pMLC and TER, as indicated, using data at respective time points shown in Figs. 1g,1d and 2a. ns=not significant.

### Dynamic regulation of VE-cadherin by Arp2/3 complex controlled actin driven JAIL in TNF-α-induced inflammatory morphological phenotype

Following the termination of the intermediate state, endothelial dynamics progressively increased, characterised by rapid MLC-rephosphorylation (Figure 2a), junctional actin disassembly, stress fibre formation (Figure 1e, 1f, and 2d), and the reappearance and gradual expansion of membrane protrusions (Figure 1e and 2c, 50-400 minutes), consistent with increased junctional dynamics and progressive shape changes. Next, we investigated the role of VE-cadherin, a key adhesion protein at endothelial junctions, in TNF-α-induced morphological remodelling, dynamics and the regulation of barrier function, with particular focus on JAIL. JAIL are specialised, actin-driven membrane protrusions, regulated by the Arp2/3 complex, that form locally at VE-cadherin-depleted junctions and directly establish new VE-cadherin adhesions at these sites. This process drives VE-cadherin dynamics and has previously been observed in angiogenesis and wound healing (54, 56, 78). Fluorescence live cell imaging was performed on HUVEC expressing moderate levels of VE-cadherin-mCherry and EGFP-p20. The use of EGFP-tagged p20, a fluorescently labelled subunit of the Arp2/3 complex, proved highly effective in visualising JAIL activity. Under control conditions, moderate JAIL dynamics were observed, characterised by the formation of moderately sized and curved VE-cadherin plaques driven by EGFP-p20-labelled Arp2/3 activity, which promotes the development of branched actin filaments. (t = 00:00 hh:mm, Figure 3a; overview is shown in Supplementary Fig. S2a). Consistent with TNF-α-induced actin dynamics, two distinct dynamic states of VE-cadherin and EGFP-p20 were observed after TNF-α stimulation.

**Figure 3.**
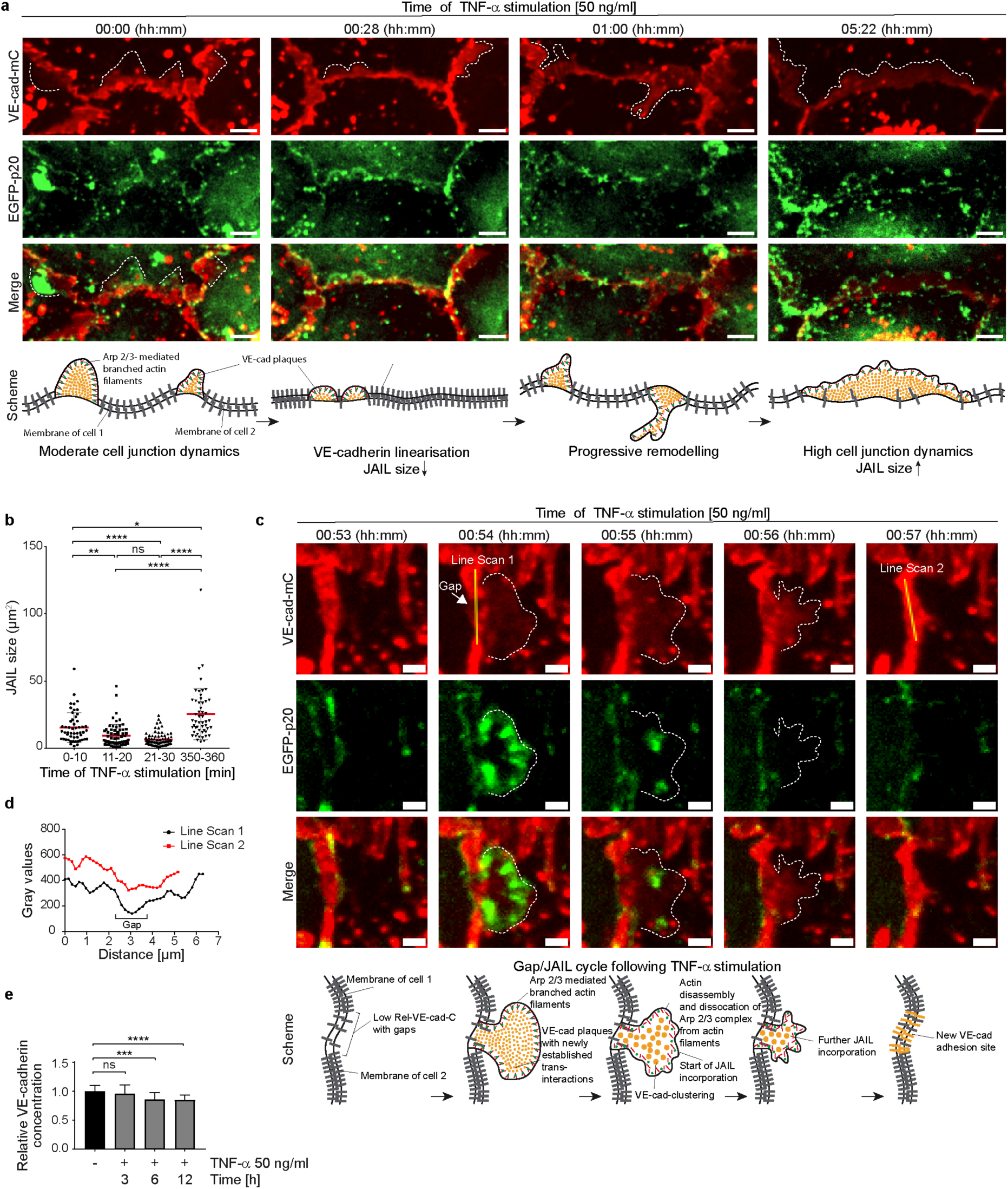
TNF-α-induced Arp2/3 complex regulates VE-cadherin dynamics through JAIL formation in intermediate and mature inflammatory phenotypes. **a)** VE-cadherin-mCherry and EGFP-p20 expressing HUVEC treated with 50ng/ml TNF-α at time points as indicated. Dotted lines indicate JAIL. EGFP-p20 is only seen during the JAIL extension. Note: Shortly after TNF-α treatment VE-cadherin undergoes linearisation with reduction in JAIL size followed by progressive increased cell junction dynamics after about 6h. Scale bar, 5 µm. **b)** Transient downregulation of JAIL size during VE-cadherin linearisation upon TNF-α stimulation. n= 3 independent experiments with t= 0-10 min, 52 JAIL; t= 11-20 min, 66 JAIL; t= 21-30 min, 75 JAIL; t= 350-360 min, 52 JAIL; Kruskal-Wallis-test. **c)** Gap/JAIL cycle depicted at approximately 1h after TNF-α stimulation during the autonomous activation phase. Scale bar, 2 µm. **d)** Line scans 1 and 2 depicted in c indicate gap and gap closure by JAIL mediated VE-cadherin dynamics. **e)** Quantification of the relative VE-cadherin concentration (intensity of VE-cadherin/given cell border length) over 12 hours upon TNF-α stimulation; n = 4 independent experiments of LSM images at 63x-maginifaction. NT, n=22; TNF-α 3h, n= 22; TNF-α 6h, n=22; TNF-α 12h, n=25. Ordinary one-way ANOVA. ns = not significant.

This included the initial intermediate state, characterised by the linearisation of VE-cadherin-mCherry at cell junctions within the first 30 minutes (Figure 3a, t = 00:28 min), accompanied by a reduction in the size of VE-cadherin plaques, indicating reduced JAIL formation and Arp2/3 activity (Figures 3a, 3b). Subsequently, the second state of VE-cadherin dynamics reversed the intermediate state. Specifically, we found a progressive upregulation of JAIL-mediated VE-cadherin dynamics (Figure 3a, 3b) within 6 hours, as evidenced by increased VE-cadherin plaque size of up to 66%. This process also contributes to endothelial shape change (Movie 2). This state represents the fully developed IMP driven by JAIL formation (Figure 3a, 3b).

To document the details of TNF-α-induced JAIL-driven VE-cadherin dynamics, a region in HUVEC cultures moderately expressing VE-cadherin-mCherry and EGFP-p20 was selected approximately 1 hour after TNF-α stimulation, capturing the ongoing remodelling process (Figure 3c, 3d). Notably, the Arp2/3 complex was only visible during the transient extension phase, whereas VE-cadherin persisted throughout the time-lapse recordings. At approximately 54 minutes, EGFP-p20-positive protrusions formed locally at junctions depleted of VE-cadherin, indicating JAIL formation. These protrusions bordered VE-cadherin plaques and rapidly disappeared after maximal expansion, while VE-cadherin plaques remained intact. Subsequently, VE-cadherin clusters formed within the plaques and integrated into junctions, confirming that TNF-α-induced VE-cadherin dynamics are driven by JAIL both during development and after establishment of the IMP. The overall time course of TNF-α-induced JAIL formation is consistent with Rac-1-mediated Arp2/3 complex activation, which is known to initiate branched actin-driven membrane protrusions that overlap adjacent cell membranes, allowing the formation of VE-cadherin trans-interactions that manifest as VE-cadherin plaques (56).

To better understand the shift from the intermediate state to the progressive increase in JAIL-mediated VE-cadherin dynamics and the impact of shape change in this process, the relative VE-cadherin concentration (Rel-VE-cad-C) (56) was determined at specific time points from time-lapse recordings after TNF-α stimulation. The Rel-VEcad-C was determined by calculating the intensity of VE-cadherin along a given junction length in relation to that junction length (56). Indeed, alongside the TNF-α-induced changes in cell shape, a progressive reduction Rel-VE-cad-C is observed, reaching an average reduction of approximately 15% after 6 hours (Figure 3e). This dilution of VE-cadherin is sufficient to enhance Rac-1-activated, Arp2/3 complex-controlled, actin-driven JAIL formation, which facilitates the local re-establishment of VE-cadherin plaques, indicating the sensitivity of this parameter. Thus, the interplay between shape changes, VE-cadherin dilution and JAIL formation is proposed as the underlying mechanism driving TNF-α-induced junctional dynamics. Since JAIL also locally restores VE-cadherin adhesion, it serves as a regulatory element for cell junctions, even within the TNF-α-induced inflammatory morphological phenotype. The properties of JAIL highlight their dual role as a key factor responsible for shape changes due to increased cell and junctional dynamics, as well as a control mechanism for preserving barrier integrity in the fully developed IMP. The entire process can be followed in movie 2.

### TNF-α-induced inflammatory morphological phenotype is driven and controlled by gap/JAIL cycles in a VE-cadherin-dependent manner

So far, we have shown that the TNF-α-induced fully developed IMP is driven by actin and VE-cadherin dynamics (JAIL), passing through a morphological intermediate state and involving a characteristic sequence of changes in barrier function and junctional dynamics. Importantly, TNF-α-induced VE-cadherin dilution and ongoing dynamics led to the transient formation of intercellular gaps (Figure 4a; Supplementary Figure S3a), which in turn stimulated the formation of JAIL, which locally restored VE-cadherin-mediated cell adhesion. These alternating and functionally coupled mechanisms therefore led to gap/JAIL cycles, which have two important effects on the endothelium. On the one hand, gap/JAIL cycles provide a mode of regulation of intercellular junctions independent of the inflammatory morphological phenotype, thereby maintaining a basal barrier function. On the other hand, the temporarily formed intercellular gaps provide exit points for neutrophil TEM.

**Figure 4.**
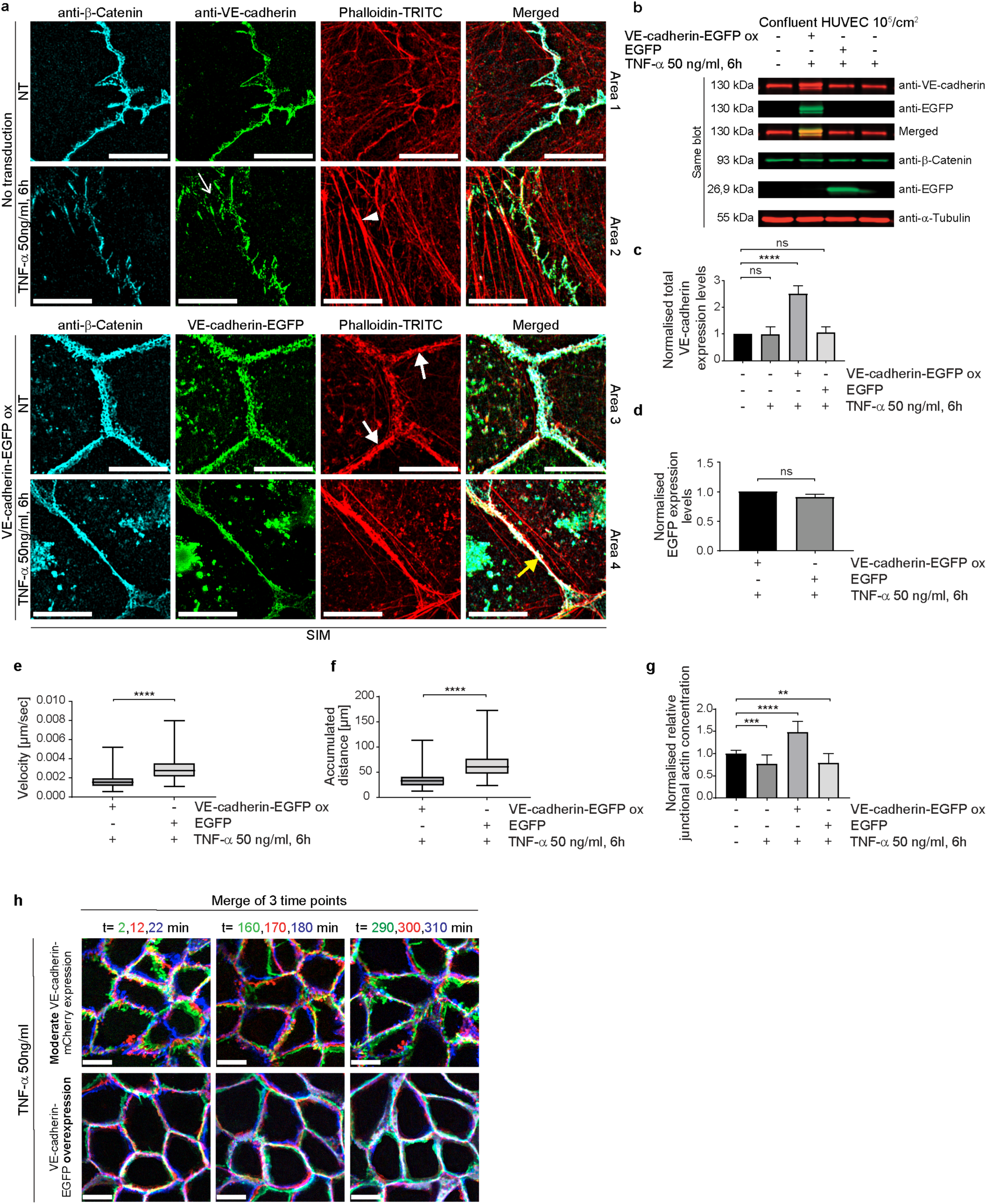
VE-cadherin-EGFP overexpression potentiates junctional actin recruitment and reduces cell-junction dynamics. **a)** SIM of TNF-α-treated control and VE-cadherin overexpressing HUVEC cultures. Cell cultures were labelled as indicated. For overviews, see Supplementary Figure S3. Upper panel, control cultures (NT) and TNF-α treated cultures as indicated. Small white arrow indicates intercellular gaps. White arrowheads indicate stress fibres. Lower panel. VE-cadherin-EGFP overexpression recruited actin to cell junctions (white bold arrow) together with β-catenin. TNF-α treated cultures still show actin recruitment colocalised with β-catenin and VE-cadherin-EGFP (yellow bold arrows). Scale bar: 10 µm. **b)** Western blot showing the expression of VE-cadherin, VE-cadherin-EGFP, β-catenin, α-tubulin (loading control) and EGFP (transfection control) in HUVEC, 6 hours after TNF-α treatment, and in controls as indicated. **c)** Quantification of VE-cadherin expression levels. 4 WBs of n = 3 independent experiments. Ordinary one-way ANOVA. **d**) Quantification of EGFP expression levels. N= 3 independent experiments of 5 Western Blots. Unpaired student t-Test. **e,f)** Dynamics of TNF-α-treated HUVEC cultures overexpressing VE-cadherin-EGFP versus EGFP-only as controls. Analyses include **e)** migration velocity, **f)** accumulated distance travelled. **e, f)** n= 3 independent experiments with analyses of 108 cells in control and 144 cells in VE-cadherin-EGFP overexpressing cultures. Mann-Whitney-U-test. **g)** Relative junctional actin recruitment in HUVEC after VE-cadherin labelling. N= 3 independent experiments with NT (n=21), TNF-α treated (n=25), VE-cadherin-EGFP overexpressing + TNF-α (n=16). **h)** VE-cadherin-mCherry displacements in HUVEC during 6 hours of TNF-α treatment with false-colour analysis at three time points with 10-minute intervals. Colours of βt =2 min, green; βt=12 min, red; βt=22 min, blue. Scale bar: 20 µm. Overexpression of VE-cadherin reduces displacement. ns = not significant.

To evaluate the proposed important role of VE-cadherin dilution towards an endothelial IMP with gap/JAIL cycles and its impact on neutrophil TEM, EGFP-labelled VE-cadherin (Ad-VEcad-EGFP) was overexpressed by adenovirus-mediated gene transfer. Ad-VEcad-EGFP (59) is able to rapidly upregulate VE-cadherin expression in HUVEC to a 2.5-fold higher level within hours (Figure 4a-c), while overexpression of EGFP alone served as a control at a comparable level (Figure 4d). The TNF-α-induced ICAM-1 expression was maintained across all experimental conditions including after overexpression of Ad-VEcad-EGFP and EGFP alone (Supplementary Fig. S1a, b). The functionality of Ad-VEcad-EGFP was further confirmed by its ability to recruit β-catenin to cell junctions (Supplementary Fig. S3a, b, c), while the total β-catenin expression tended to increase but remained insignificant (Supplementary Fig. S3d). Enrichment of Ad-VEcad-EGFP along the cell borders reduced stress fibres (Figure 4a; Supplementary Fig. S3a), downregulated cell migration even after TNF-α treatment (Figures 4e, f; Supplementary Fig. S3e) and resulted in approximately 50% greater recruitment of junctional actin filaments compared to control cells (Figure 4g). Furthermore, TNF-α treatment altered neither the VE-cadherin level nor junctional actin recruitment (Figure 4c, g), indicating the important role of VE-cadherin in the development of an IMP. This is further underlined by the fact that VE-cadherin overexpression not only blocked cell migration, but was also associated with a general down regulation of cell and cell junctional dynamics (Figures 4e, f; Supplementary Fig. S3e, Movie 1). The inhibition of the overall morphodynamics in Ad-VEcad-EGFP-overexpressing cells is attributed to increased clustering of VE-cadherin along junctions and strong recruitment of junctional actin, which likely interacts with VE-cadherin and prevents its lateral displacement, as demonstrated by false colour analysis in time-lapse recordings (Figure 4h; Movie 1, right panel). The data also show that increased VE-cadherin clustering enhances actin recruitment, highlighting a time-dependent sequence in which VE-cadherin clustering precedes actin recruitment. Accordingly, lateral displacement of VE-cadherin appears to be critical for TNF-α signalling, leading to disassembly of junctional actin and weakening of cell junctions to facilitate remodelling as this was inhibited by VE-cadherin overexpression. Furthermore, VE-cadherin overexpression prevents TNF-α-induced development of the IMP and gap/JAIL cycles, whereas TNF-α-induced ICAM-1 expression occurs independently of VE-cadherin expression, suggesting different TNF-α mediated signalling pathways in this process towards a fully developed inflammatory phenotype.

### Impaired TNF-α-induced junctional dynamics via VE-cadherin overexpression blocks neutrophil transmigration

To characterise the effect of gap/JAIL cycles on neutrophil TEM, HUVEC were exposed to TNF-α for 6 hours to allow the development of the IMP and TEM was assessed under low shear stress conditions of 1 dyn/cm². Shear stress was generated using a custom-designed parallel plate flow chamber (insert flow chamber, patent number: 102020131849), (Supplementary Fig. S4a). Neutrophils introduced into the medium-stream via a side port exhibited characteristic margination/capture, rolling, firm adhesion, spreading and crawling on the surface of TNF-α-activated control HUVEC, followed by transendothelial migration (TEM) (Movie 3, left panel). Overexpression of Ad-VEcad-EGFP in HUVEC displayed an undisturbed neutrophil adhesion to the TNF-α activated cells (Figure 5a, Supplementary Movie 3, right panel). In contrast, we found a reduced neutrophil TEM by approximately 3-fold (80%) (Figure 5b). Live cell imaging (Movie 4) showed that neutrophils probe the VE-cadherin-overexpressing endothelial cell layer for suitable transmigration sites, while TEM is reduced. Calculation of the number of adherent neutrophils in relation to the number of transmigrated neutrophils confirms this observation. Overexpression of EGFP alone as a further control has neither an effect on TEM rates or adhesion compared to untreated controls (Figure 5a, 5b).

**Figure 5.**
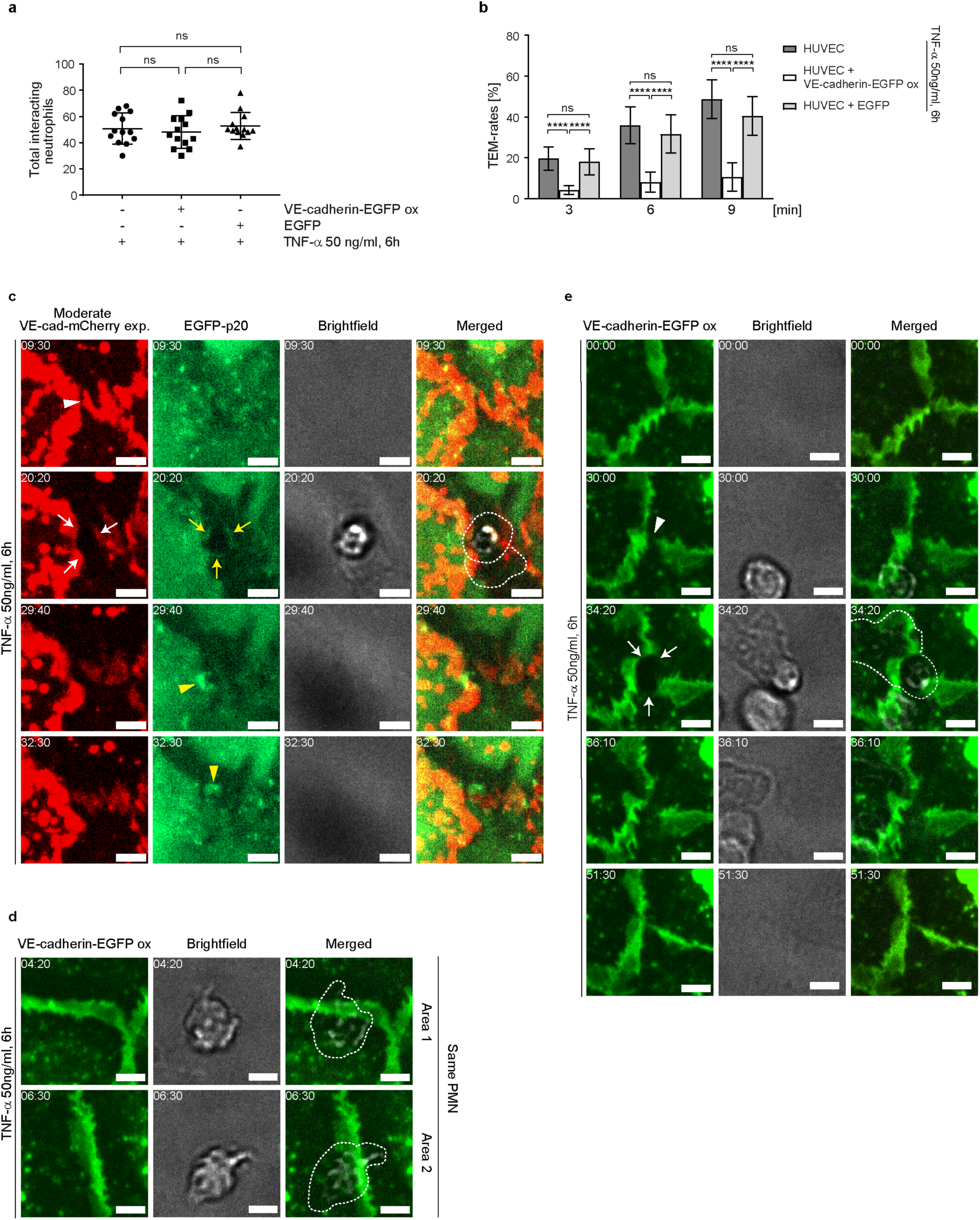
VE-cadherin dilution and increased cell junction dynamics upon TNF-α treatment are essential for neutrophil transmigration under flow. **a-b)** Neutrophil adhesion and time-dependent transmigration through TNF-α-activated native (control 1), VE-cadherin-EGFP-overexpressing and EGFP-overexpressing (control 2) HUVEC cultures placed in the “insert flow chamber” at a flow of 1 dyn/cm^2^. Analyses are based on phase-contrast time-lapse imaging (10x objective) with 10^6^ neutrophils applied to the respective HUVEC cultures. Total interacting neutrophils were counted first, followed by transmigrating neutrophils. Data represent n= 3 independent experiments for each condition, with 13 different positions analysed per experiment. **c)** Representative area showing the dynamics of moderately expressed VE-cadherin-mCherry and EGFP-p20 in TNF-α activated HUVEC cultures during neutrophil transmigration, visualised by time-lapse imaging. Intercellular gaps (white arrowheads) are used for transendothelial migration (TEM), while VE-cadherin-mCherry is displaced laterally (white arrows). Yellow arrows indicate EGFP-p20. VE-cadherin plaques document junction-associated intermittent lamellipodia (JAIL)-mediated sealing of transmigratory gaps (yellow arrowheads). Time code = mm:ss, scale bar: 5 µm. **d)** Representative area showing the dynamics of neutrophil behaviour in VE-cadherin-EGFP overexpressing HUVEC cultures. Although neutrophils show stable adhesion to intercellular spaces, transendothelial migration (TEM) was not observed. Cropped areas are taken from the time-lapse recording (Suppl. Movie 4) and as shown in Supplementary Fig. S4. Time code = mm:ss, scale bar: 5 µm. **e)** Even in VE-cadherin-EGFP overexpressing HUVEC, intercellular gaps can occur (arrowheads), which are then used by neutrophils for transendothelial migration (TEM) (arrows). Time code = mm:ss, scale bar: 5 µm. ns= not significant.

The dynamics of TEM of neutrophils through cell junctions was further assessed after moderate expression of VE-cadherin-mCherry together with EGFP-p20 to label the Arp2/3 complex as a marker of branched actin filaments. TEM of neutrophils were predominantly observed at sites where moderately expressed VE-cadherin-mCherry was reduced or absent, including intercellular gaps and tricellular junctions (Figure 5c, Supplementary Fig. S4b). Neutrophil transmigration preferentially occurred at VE-cadherin free gaps with EGFP-p20 positive docking structure like protrusions (Figure 5c), consistent with previous reports (24). After transmigration, JAIL re-established VE-cadherin adhesion and resealed the transmigration gap by forming new VE-cadherin adhesion sites (Figure 5c).

Although VE-cadherin overexpression largely blocked the TNF-α-induced development of the inflammatory morphological phenotype, a few neutrophils still transmigrated. In order to visualise the junctional conditions at these sites, we carefully searched for TEM of neutrophils in Ad-VEcad-EGFP overexpression HUVEC by life cell imaging. TEM was never observed at sites where Ad-VEcad-EGFP was densely packed, while crawling neutrophils probed the endothelium for exit points, passing several cell junctions but largely failed to find suitable sites for TEM (Figure 5d; Supplementary Fig. S4c, Movie 4 left panel). However, we were able to find occasional TEM events. Consistent with TEM events under control conditions, these few sites exhibited VE-cadherin dynamics with a local decrease in relative VE-cadherin concentration and small intercellular gaps. At these sites, TEM occurred and VE-cad-EGFP appeared to be displaced, accompanied by the formation of a transmigration pore (Figure 5e; Supplementary Fig S4d, Movie 4 right panel). These live cell imaging observations in VE-cadherin overexpressing cells confirm the requirement for intercellular gaps to allow TEM. In all cases, VE-cadherin resealed the gap after the neutrophil had passed through the cell layer, consistent with the formation of lamellipodia (44, 79) and JAIL (Figure 5e; Movie 4 right panel). In summary, the data provide compelling evidence that TNF-α-induced shape changes – leading to dilution and redistribution of VE-cadherin at endothelial junctions – together with increased junctional dynamics via gap/JAIL cycles, are defining features of the inflammatory morphological phenotype. Together with the upregulation of cell adhesion molecules, gap/JAIL cycles serve as a critical predictor for neutrophil transendothelial migration (TEM) and for the control of endothelial barrier function in inflammation. The proposed entire mechanism is illustrated below (Figure 6).

**Figure 6.**
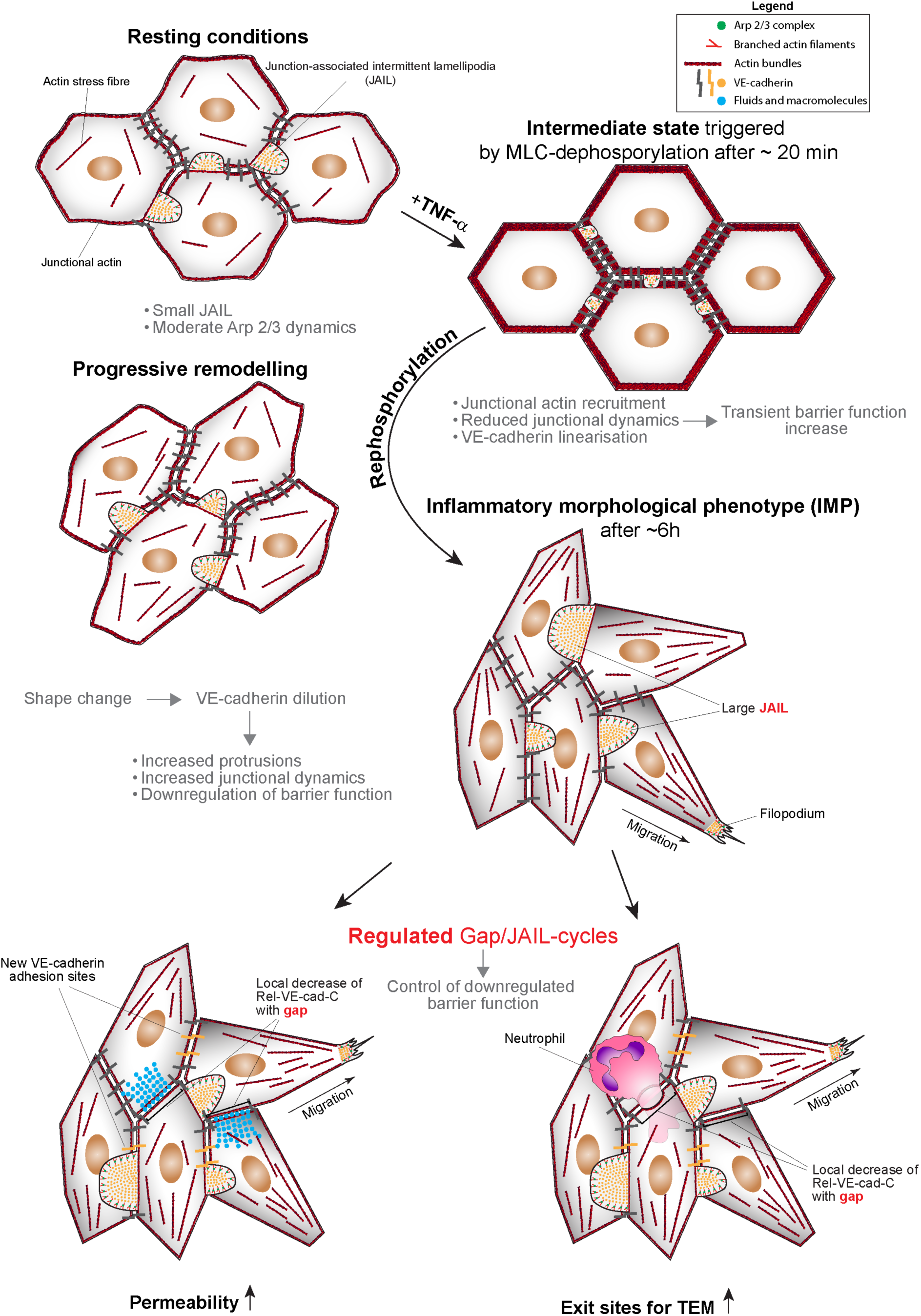
Scheme illustrating the proposed mechanistic follow up The intermediate state is characterised by junctional actin recruitment, VE-cadherin linearisation, increased barrier function and reduced cell junction activity, triggered by myosin light chain (MLC) dephosphorylation. The switch to a pro-inflammatory phenotype is then activated by autoregulated MLC rephosphorylation, which stimulates actin dynamics. This progressively leads to junctional actin disassembly, stress fibre formation, and membrane protrusions that drive Arp2/3 complex-controlled junction-associated intermittent lamellipodia (JAIL), resulting in progressive downregulation of barrier function and shape change. In particular, the shape change dilutes VE-cadherin along the junctions, reducing cell adhesion and forming intercellular gaps for neutrophil transendothelial migration (TEM), and explains the decrease in barrier function, both characteristics of the inflammatory endothelial phenotype.

## Discussion

This study investigated the TNF-α-induced inflammatory morphological phenotype (IMP) in HUVEC cultures and its effect on neutrophil transendothelial migration (TEM). TNF-α induced a biphasic morphological response: an intermediate state followed by the full development of the IMP. The intermediate state, characterised by MLC dephosphorylation, junctional actin assembly, VE-cadherin linearisation and transient barrier enhancement, may act as a gatekeeper to prevent monolayer collapse and uncontrolled leakage. This state resembles quiescent endothelium and may represent a reset point for transition to the IMP. Subsequent rephosphorylation of MLC induced stress fibre formation, junctional actin disassembly and VE-cadherin dilution, resulting in intercellular gaps and junction-associated intermittent lamellipodia (JAIL), forming gap/JAIL cycles essential for regulating barrier function and neutrophil TEM.

### The TNF-α-induced intermediate state: A gatekeeper of inflammation

Intermediate cell states occur in various cellular processes such as epithelial-mesenchymal transitions and developmental biology, and are supposed to play an important role in coordinating these processes (80, 81). The intermediate state following TNF-α prevents the monolayer from collapse with uncontrolled leakage of fluid and macromolecules. This is a reasonable assumption, as the intermediate state is characterised by rapid junctional actin recruitment, VE-cadherin linearisation and downregulated morphodynamics/junctional dynamics, leading to a transient increase in barrier function, consistent with previous reports (82). The TNF-α-induced intermediate cell state resulted from the dephosphorylation of myosin light chain (MLC), a critical regulator of barrier function and contractility (76, 83). This intermediate state may be further involved in the induction of genetic and epigenetic regulation after TNF-α stimulation as described (84, 85). However, this needs to be evaluated separately.

### The role of VE-cadherin dynamics and gap/JAIL cycles in the TNF-α induced inflammatory phenotype

The full development of the TNF-α-induced inflammatory endothelial phenotype is characterised by the expression of cell adhesion molecules and by the morphological transformation leading to gap/JAIL cycles. The development of the IMP is mediated by two key mechanisms. First, VE-cadherin dilution occurred as cell shape changes (e.g., elongation) geometrically elongated junctions, while compensatory pathways maintained overall expression levels despite TNF-α-driven VE-cadherin degradation (86). Second, TNF-α-induced VE-cadherin dilution and intercellular gaps stimulate an increase in actin-driven JAIL formation regulated by Arp2/3 complex activity to dynamically restore VE-cadherin adhesion at these sites, in agreement with observation in angiogenesis and wound healing (for review see (56)). As a result of VE-cadherin dilution, intercellular gap formation and JAIL-mediated VE-cadherin remodelling driving gap/JAIL cycles occur that continue to control junctional integrity independent of the inflammatory morphological phenotype. The transition from the intermediate state to the fully developed IMP involves MLC rephosphorylation-dependent loss of junctional actin, stress fibre formation and increased actin-driven JAIL formation. Gap/JAIL cycles balance barrier integrity and permeability during inflammation and function as a fundamental mechanism in cell junction dynamics.

### The role of VE-cadherin dilution in neutrophil transmigration

TNF-α-induced gap/JAIL cycles determine cell junction dynamics and characterise the inflammatory phenotype. Overexpression of VE-cadherin, which blocks gap/JAIL cycles, reduces TEM by ∼80% while CAM expression is unaffected, which is in agreement with previous studies showing VE-cadherin-independent regulation of ICAM-1 (Bui et al., 2020). The inhibition of TEM by VE-cadherin overexpression highlights the critical role of VE-cadherin dilution and VE-cadherin-depleted gaps forming gap/JAIL cycles for TEM. Although gap/JAIL cycles are indispensable for neutrophil TEM, particularly to define the specific exit sites for TEM, the subsequent diapedesis requires a coordinated interplay of adhesion molecules and junctional proteins, including ICAM-1 (87–89), endothelial-specific signals (90, 91), PECAM-1 and CD99 among others (17, 92). These data are consistent with the ‘path of least resistance’ concept (45), whereby increased cortical actin and VE-cadherin linearisation reduced paracellular TEM. Established research regarding VE-cadherin internalisation (37, 93), membrane recycling (32) and VE-cadherin stabilisation (2, 6, 17, 89, 94, 95) further supports the concept that resistance reduction is required along the route of TEM, underscoring the essential role of VE-cadherin in this process. Following completion of TEM, transmigratory gaps are closed via ARP2/3 complex-driven membrane protrusions (24), which are thought to reestablish VE-cadherin adhesion. Our data directly demonstrate this process, supporting its consistency with JAIL formation, whereas others have proposed that such membrane protrusions may trap leukocytes to facilitate TEM (96). Finally, VE-cadherin dilution along junctions facilitates neutrophil transmigration but is also modulated by platelets, inflammatory mediators and proteases (16, 97–100).

### Conclusion and outlook

In conclusion, this study extends the understanding of neutrophil transmigration through TNF-α activated endothelial monolayers based on gap/JAIL cycles in addition to the well-known expression of cell adhesion molecules for this process. The development of the endothelial IMP requires a complex dynamic interplay between actin, VE-cadherin, intercellular gaps and barrier function leading to gap/JAIL cycles. The identification of an intermediate state prior to IMP formation reveals a previously unknown phenomenon that enhances our understanding of endothelial cell biology in inflammatory contexts. While the full functional significance of this state – particularly in terms of its genetic and epigenetic regulation – remains to be elucidated, its primary effect is likely to be a protective mechanism that stabilises cell junctions, thereby preventing rapid junctional disassembly in response to TNF-α, and may allow the coordination of hierarchical development of the IMP.

However, the primary finding of this study demonstrates that shape change-mediated VE-cadherin dilution, and its subsequent role in driving gap/JAIL cycling, critically regulates endothelial barrier integrity and neutrophil transendothelial migration (TEM) during inflammatory responses. Pharmacological control of gap/JAIL cycles – or even one of them – may have important implications for the development of novel therapeutic strategies aimed at modulating inflammation and preventing excessive leukocyte extravasation in various disease states. Modulation of gap/JAIL cycle dynamics may also provide a means to fine-tune the inflammatory response and protect against endothelial dysfunction. Further research will elucidate the detailed molecular mechanisms underlying these processes and validate them in in vivo models to explore their therapeutic potential. The entire mechanisms and phenomena are illustrated in figure six (Figure 6).

## Acknowledgements

We thank Annelie Ahle, Franziska Merten and Christine Schimp for their skillful technical assistance. Masatoshi Takeichi generously provided us with VE-cadherin-EGFP adenoviral constructs, and Milos Galic provided expert advice on image analysis. We are very grateful to Till Rauterberg for his editing and help in preparing the movies. Special thanks to Dietmar Vestweber for critical discussion and expertise on leukocyte transmigration. The work was supported by grants from the Deutsche Forschungsgemeinschaft (DFG) and the BMBF (03ZZ0902D) to H.S. (SCHN 430/9-1 and DFG INST 2105/24-1) and from the Max Planck Society M.O-S, H.S. We are grateful for the generous support of Werner Paulus and colleagues at the Institute of Neuropathology, University Hospital Münster.

## Author contributions

Author contributions J.P.K. performed the majority of the experiments, analysed the data and prepared the figures with significant help of M.O-S. who also performed the phosphorylation and impedance spectroscopy measurements and assisted in designing and optimizing of the “Insert Flow Chamber”. V.B. contributed to overexpression experiments together with J.F. who also helped with the movie preparation. J.S assisted in analyses of cell morphologies. M.A. performed LifeACT-EGFP experiments. H.S. initiated the topic, supervised the entire work, and wrote the manuscript together with J.P.K. All authors contributed with discussions, suggestions and critical reading and editing the MS.

---

**Supplementary Figure S1.**
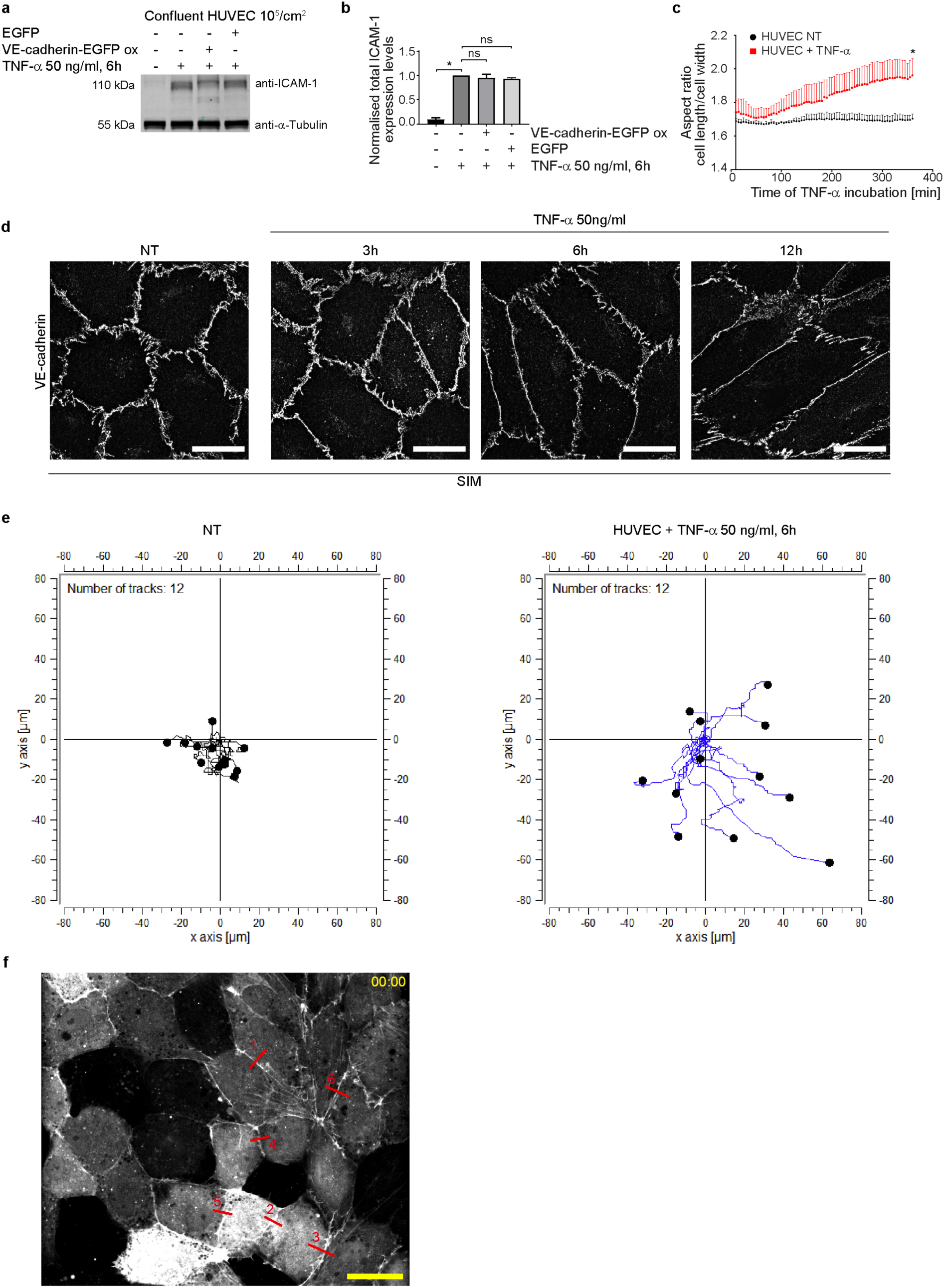
Western blot analysis of ICAM-1 from naive, EGFP– and VE-cadherin-EGFP-overexpressing HUVEC after 6 hours of TNF-α treatment. **a)** Western blot analyses of HUVEC treated as indicated. **b**) Quantification of ICAM-1 expression levels (a,b: n=3 independent experiments; Kruskal-Wallis test). **c)** Analysis of the cell aspect ratio upon TNF-α treatment; n=3 independent experiments, considering the mean values of 6 different locations at 10x magnification. Each position contains approximately 1800 +/-200 cells per time point; unpaired t-test. **d)** Shape change in HUVEC after 3h, 6h and 12h of TNF-α stimulation. Scale bar: 20 µm. **e)** Cell migration plots of control HUVEC and after TNF-α stimulation. **f)** Overview used for line scan analysis depicted in Figure 1 g-i. Scale bar 30 µm. ns= not significant.

**Supplementary Figure S2.**
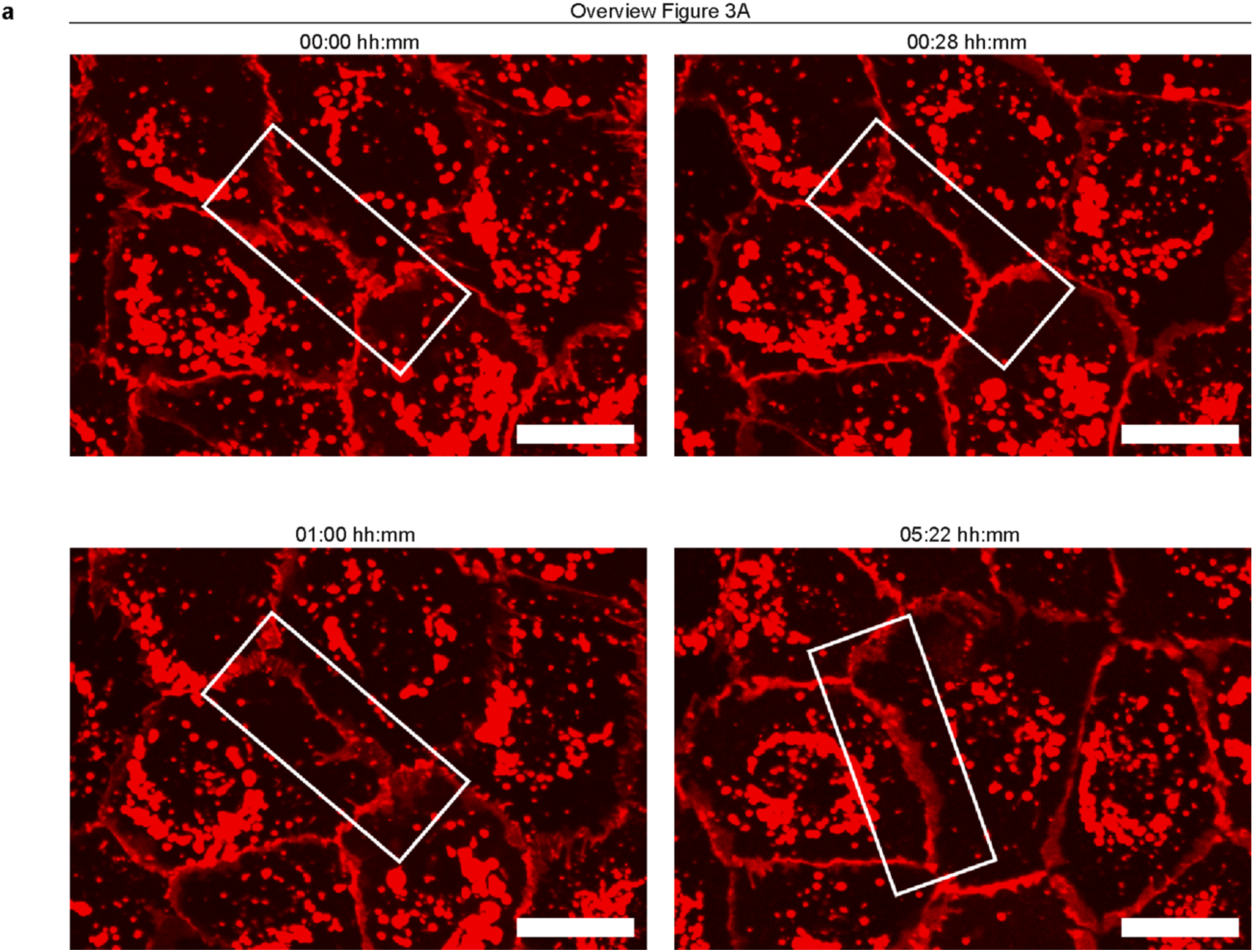
Overview of Figure 2a. Scale bar: 20 µm.

**Supplementary Figure S3.**
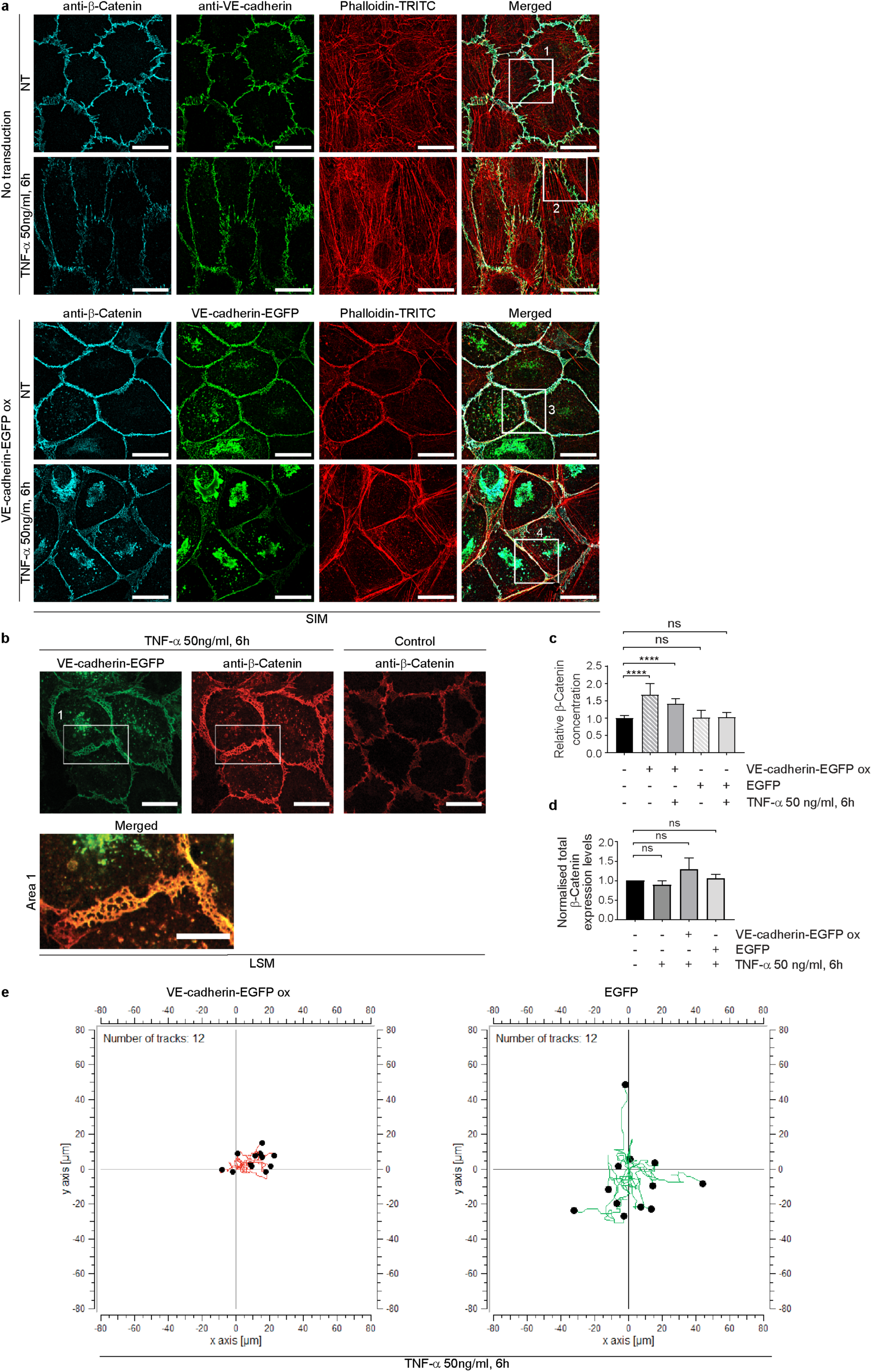
**a**) Overview of Figure 4a. **b)** LSM-images of VE-cadherin and β-Catenin. Note: Orange colour indicates VE-cadherin-EGFP and β-Catenin superimposition. **c)** Quantification of the relative β-Catenin concentrations of LSM-images as depicted in b in transduced and untransduced cells following TNF-α treatment. N= 3 independent experiments with the following number of LSM images (63x magnification) included in total per condition. NT, n= 52; VE-cadherin-EGFP, n=17; VE-cadherin-EGFP+ TNF-α, n=33; EGFP, n=18; EGFP+ TNF-α, n= 33; ordinary one-way ANOVA. Scale bar overview, 20 µm; scale bar cropped areas, 10 µm. **d)** Quantification of β-catenin expression levels, as analysed by Western blot in Figure 4b. 4 WBs from n= 3 independent experiments; Ordinary one-way ANOVA. **e)** Cell migration plots of VE-cadherin-EGFP and EGFP overexpressing HUVEC following TNF-α stimulation. 12 single cells per condition are shown. ns= not significant.

**Supplementary Figure S4.**
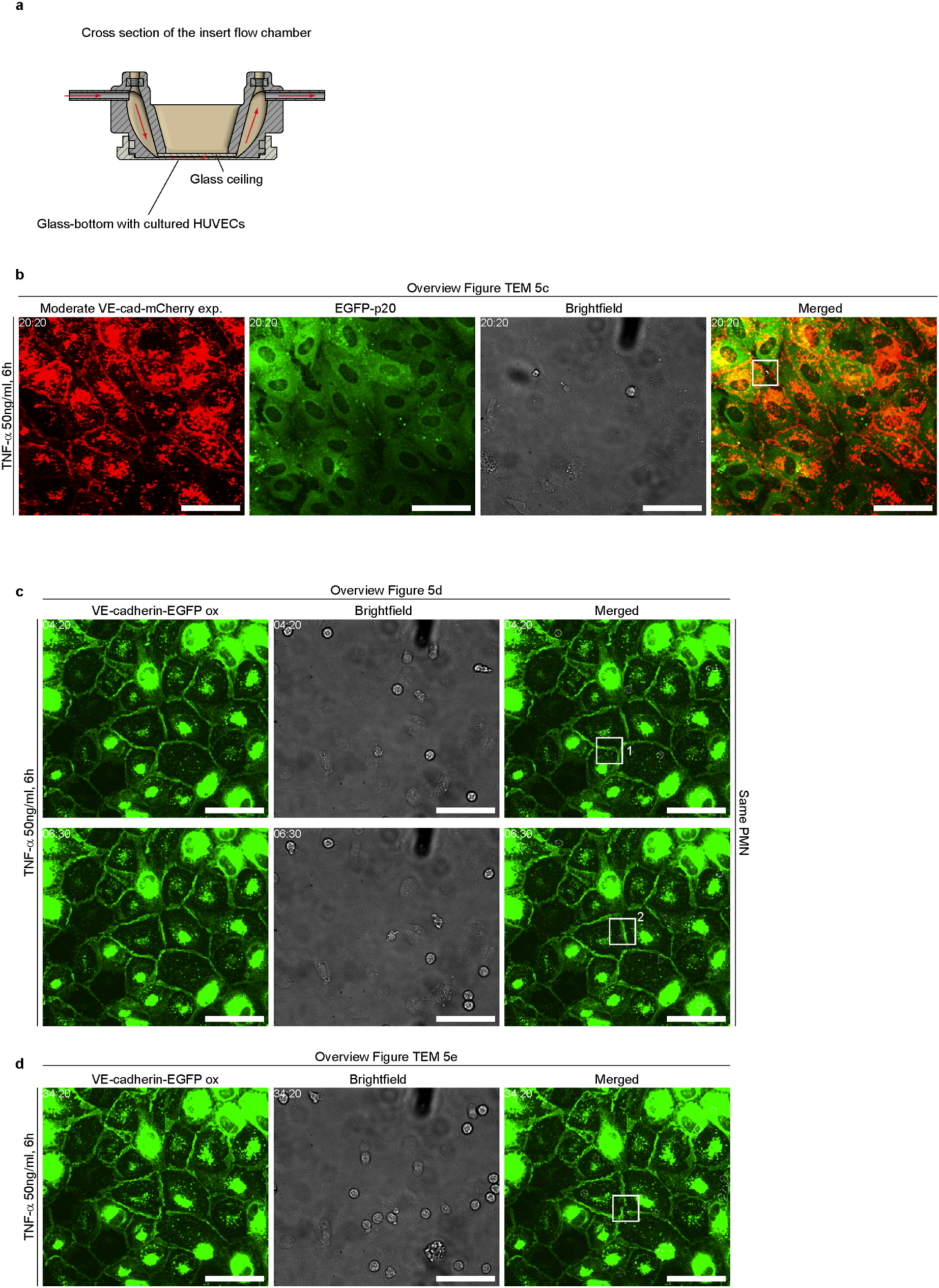
**a**) Cross section of the custom-made insert flow chamber. **b)** Overview of TEM shown in Figure 5c. Scale bar: 20 µm. **c**) Overview of two areas shown in Figure 5d. Scale bar: 20 µm. **d**) Overview of TEM depicted in Figure 5e. Scale bar: 20 µm.

